# *PIF1*, *RAD54* and *RDH54/TID1* promote residual double strand break repair during meiosis in the budding yeast, *Saccharomyces cerevisiae*

**DOI:** 10.64898/2026.01.05.697805

**Authors:** Raunak Dutta, David Murtha, Tyler Nagosky, Nancy M. Hollingsworth

## Abstract

In the budding yeast, *Saccharomyces cerevisiae*, repair of programmed double strand breaks occurs in two phases during prophase I of meiosis. During Phase 1 interhomolog recombination is mediated by the meiosis-specific Dmc1 recombinase. Crossover-specific recombination intermediates enable synapsis of homologous chromosomes, resulting in a transition to Rad51-mediated recombination in Phase 2 that repairs any residual double strand breaks so that chromosomes are intact when cells progress into the first meiotic division. Studying Phase 2 recombination is challenging because the number of breaks present at pachynema (the prophase I stage when all the homologs are synapsed) is small and a low frequency of new breaks continues to be made. Using a newly developed method for analyzing Phase 2 recombination, this work discovered that *RDH54*/*TID1* can partially compensate for *RAD54*, while *PIF1* functions independently from both *RAD54* and *RDH54*/*TID1* in this process.

**ARTICLE SUMMARY:** Meiotic recombination involves repair of numerous programmed double strand breaks (DSBs). Failure to repair all the DSBs results in inviable gametes. Meiotic DSB repair occurs in two phases. In Phase 1, recombination occurs between homologs to create crossovers needed for proper chromosome segregation. Subsequently, Phase 2 recombination repairs any remaining DSBs prior to cells progressing through Meiosis I. Using a novel method for studying Phase 2 recombination, this work shows that *RDH54/TID1* can partially substitute for the related *RAD54* translocase and that the conserved *PIF1* helicase functions independently of both *RDH54/TID1* and *RAD54* in this process.

## INTRODUCTION

Meiotic recombination is a highly conserved process by which hundreds of double strand breaks (DSBs) are purposely introduced into a cell’s genome by the highly conserved Spo11 protein (KEENEY *et al*. 2014). The ends of the breaks are then resected, thereby creating single strand DNA sequences that can probe the genome for a chromosome’s homologous partner (ZICKLER AND KLECKNER 1999; HUNTER 2015; MIMITOU *et al*. 2017). A subset of these DSBs are repaired using homologs as templates to generate reciprocal crossovers that, in combination with sister cohesion, physically connect homologs, thereby enabling them to align properly on the meiosis I (MI) spindle (PETRONCZKI *et al*. 2003).

While hundreds of DSBs are needed for homolog pairing, just a single crossover is sufficient to connect two homologs. Since gametes containing even a single DSB cannot make viable progeny, it is critical that all DSBs are repaired before exiting prophase I. One solution is to have two phases of recombination. During Phase 1, DSB repair generates the requisite number of interhomolog pre-crossover intermediates necessary for chromosome segregation, with many of the remaining DSBs being repaired as non-crossovers. In Phase 2, any remaining DSBs are repaired using sister chromatids as templates. Evidence for a shift from interhomolog to intersister recombination at the mid to late pachytene transition during meiotic prophase I has been observed in the nematode, *C. elegans*, in studies using irradiated mouse spermatocytes, as well as in the budding yeast, *Saccharomyces cerevisiae* (COLAIACOVO *et al*. 2003; HAYASHI *et al*. 2007; ARGUNHAN *et al*. 2017; PRUGAR *et al*. 2017; ENGUITA-MARRUEDO *et al*. 2019; TORAASON *et al*. 2021).

The transition from Phase 1 to Phase 2 recombination in budding yeast is controlled by the meiosis-specific kinase, Mek1 (ROCKMILL AND ROEDER 1991; LEEM AND OGAWA 1992; CHEN *et al*. 2018). Mek1 regulates several steps of recombination, including DSB end resection, interhomolog bias, and the crossover-specific ZMM pathway (KIM *et al*. 2010; CHEN *et al*. 2015; KRYSTOSEK AND BISHOP 2024). In addition, Mek1 is the effector kinase for the meiotic recombination checkpoint (MRC) that prevents cells from exiting prophase I when a threshold number of DSBs are present (LYDALL *et al*. 1996; XU *et al*. 1997; MACQUEEN AND HOCHWAGEN 2011; CHEN *et al*. 2018). The checkpoint kinases, Mec1 and Tel1 are recruited to DSBs where they phosphorylate the meiosis-specific chromosome axis protein Hop1 (HOLLINGSWORTH *et al*. 1990; SMITH AND ROEDER 1997; CARBALLO *et al*. 2008; CHUANG *et al*. 2012). Mek1 binds to phosphorylated Hop1 through its forkhead associated (FHA) domain and autoactivates by phosphorylation *in trans* (NIU *et al*. 2007). Mek1 kinase activity therefore provides an indirect readout for DSBs (PRUGAR *et al*. 2017). When there is a sufficient number of DSBs, Mek1 phosphorylates the DNA binding domain of the meiosis-specific transcription factor, Ndt80, thereby preventing the expression of >300 genes, including those needed to disassemble the SC and exit from prophase I (CHU AND HERSKOWITZ 1998; SOURIRAJAN AND LICHTEN 2008; CHEN *et al*. 2018). Mek1 inhibition of Ndt80 therefore keeps cells in prophase I when there is a threshold number of DSBs, raising the question how does a cell know when sufficient DSBs have been processed into pre-crossover intermediates so that any remaining breaks can be repaired, allowing the cell to progress to metaphase I?

In many organisms, including yeast and mammals, the 3’ ends of resected DSBs are bound by two recombinases, Rad51 and the meiosis-specific Dmc1, to form nucleoprotein presynaptic filaments that search the genome for homology and then invade the homologous sequence to create displacement or D-loops (BROWN AND BISHOP 2014). Rad51 strand exchange activity is not required for meiotic interhomolog recombination, but the presence of the Rad51 protein on the presynaptic filament is necessary (CLOUD *et al*. 2012; DA INES *et al*. 2013). In mitotically dividing cells and *in vitro*, Rad51 interacts with Rad54, is a Swi2/Snf2 double-stranded DNA translocase, to make D-loops (CLEVER *et al*. 1997; GOLUB *et al*. 1997; PETUKHOVA *et al*. 1998; PETUKHOVA *et al*. 1999; HEYER *et al*. 2006; WRIGHT AND HEYER 2014; CRICKARD 2021). However in meiotic yeast cells, Rad51-Rad54 interaction is prevented during Phase 1 by Mek1 phosphorylation of both Rad54 and a meiosis-specific protein, Hed1, that inhibits Rad51-Rad54 complex formation (TSUBOUCHI AND ROEDER 2006; BUSYGINA *et al*. 2008; NIU *et al*. 2009; CALLENDER *et al*. 2016). As a result, Dmc1, in combination with the Rad54 paralog, Rdh54/Tid1, is the sole recombinase able to mediate interhomolog strand invasion when Mek1 activity is high (DRESSER *et al*. 1997; SHINOHARA *et al*. 1997; ARBEL *et al*. 1999; CHI *et al*. 2009; NIMONKAR *et al*. 2012). (To prevent confusion between Rad54 and Rdh54, Tid1 will be used hereafter to designate Rdh54). Phase 1 recombination is defined as the Dmc1-driven interhomolog DSB repair that occurs in prophase I when Mek1 kinase activity is high (ZIESEL *et al*. 2022).

Processing of D-loops by a specific set of meiosis-specific proteins called the ZMMs (Zip1, Zip2, Zip3, Zip4, Msh4, Msh5, Mer3, Spo16), creates crossover precursor intermediates that promote chromosome synapsis (ALLERS AND LICHTEN 2001; BÖRNER *et al*. 2004; LYNN *et al*. 2007; PYATNITSKAYA *et al*. 2019; PYATNITSKAYA *et al*. 2022). The synaptonemal complex (SC) is a tri-partite protein structure formed when homologs synapse (PAGE AND HAWLEY 2004). In an SC from yeast, pairs of condensed sister chromatids are connected by a central region containing the transverse filament protein, Zip1 and the central element proteins Ecm11 and Gmc2 (SYM *et al*. 1993; HUMPHRYES *et al*. 2013). Pachynema is the prophase I stage during which homologs are fully synapsed. Insertion of the central region removes the bulk of the DSB machinery and Mek1 from the axes, thereby decreasing both DSB formation and Mek1 kinase activity (SUBRAMANIAN *et al*. 2016; PRUGAR *et al*. 2017; MU *et al*. 2020; LEE *et al*. 2021). As a result, Ndt80 activates transcription of many genes, including the polo-like kinase Cdc5 (CHU AND HERSKOWITZ 1998). Cdc5 activity triggers resolution of recombination intermediates into crossovers, degradation of Red1 resulting in complete inactivation of Mek1 and exit from prophase I (SOURIRAJAN AND LICHTEN 2008; PRUGAR *et al*. 2017; CHEN *et al*. 2018). Coupling synapsis, which only occurs when DSBs are repaired in a way that generates pre-crossover intermediates, to Mek1 regulation of Ndt80 ensures that cells progress into MI with physically connected homologs.

Phase 2 recombination occurs in pachytene cells to repair any remaining DSBs. Although synapsis greatly reduces the frequency of Spo11 DSBs, break formation is not completely eliminated, with ∼0-10 DSBs per pachytene nucleus (XU *et al*. 1995; ALLERS AND LICHTEN 2001; THACKER *et al*. 2014; SUBRAMANIAN *et al*. 2016). These DSBs are not needed to make crossovers but must still be repaired so that chromosomes are intact at MI. Several experiments support the idea that Phase 2 recombination utilizes Rad51-Rad54. (1) Deletion of *RAD54* has no effect on interhomolog recombination, yet reduces spore viability to ∼50% (SHINOHARA *et al*. 1997). Similarly, a strand exchange-specific mutant of Rad51, *rad51-II3A*, exhibits wild-type Phase 1 recombination, but produces ∼80% viable spores (CLOUD *et al*. 2012). This reduced spore viability is not due to MI non-disjunction, as is the case for mutants that fail to make interhomolog crossovers (ZIESEL *et al*. 2022). (2) Rad51-Rad54 interaction is regulated by Mek1 phosphorylation, a reversible modification that can be removed by phosphatases when Mek1 activity goes down at pachynema (NIU *et al*. 2009; CALLENDER *et al*. 2016). (3) *NDT80* transcription can be artificially controlled using an *NDT80* inducible system (*NDT80-IN* for short). This system combines *P_GAL1_-NDT80* with a gene that fuses the *GAL4* transcriptional activator to a fragment of the human estradiol receptor (*GAL4-ER*) (BENJAMIN *et al*. 2003; PRUGAR *et al*. 2017). Addition of estradiol (ED) activates transcription of *P_GAL1_-NDT80*. When cells are incubated in Sporulation (Spo) medium in the absence of ED, they arrest in pachynema, similar to an *ndt80Δ* diploid (XU *et al*. 1995; BENJAMIN *et al*. 2003). Addition of ED allows cells to complete recombination, exit pachynema, progress through the meiotic divisions and sporulate. The spore viability defect of an *NDT80-IN rad54Δ* diploid can be rescued by co-induction of *RAD54* with *NDT80-IN*, showing that *RAD54* functions after Phase 1 (PRUGAR *et al*. 2017). (4) When Rad54 is depleted from the nuclei of *ndt80Δ*-arrested cells, there is a dramatic increase in Rad51 foci, a cytological indicator of DSBs (BISHOP 1994; SUBRAMANIAN *et al*. 2016). This phenotype is not observed when Rad54 is co-depleted with Mer2, a protein required for meiotic DSB formation, confirming that the Rad51 foci in the Rad54-depleted diploid arise from Spo11 generated breaks (ROCKMILL *et al*. 1995; SUBRAMANIAN *et al*. 2016). (5) At some, but not all, DSB hotspots, interhomolog bias is decreased as expected due to the reduction in Mek1 activity (SUBRAMANIAN *et al*. 2016). These results suggest there is a shift from Dmc1/Tid1-mediated interhomolog recombination during Phase 1 to Rad51/Rad54 mediated recombination in pachynema in Phase 2.

Pif1 is a 5’-3’ DNA helicase that promotes genome stability in a variety of ways, including regulation of telomerase length, Okazaki fragment processing, and replication through the ribosomal DNA (BYRD AND RANEY 2017). In addition, Pif1, in conjunction with PCNA, promotes Polymerase δ extension of Rad51-mediated D-loops during break induced replication in vegetative cells (SAINI *et al*. 2013; WILSON *et al*. 2013; BUZOVETSKY *et al*. 2017). Pif1 activity is inhibited during Phase 1 meiotic recombination and therefore is not required for processing of Dmc1-generated D-loops (VERNEKAR *et al*. 2021). However *PIF1* promotes Phase 1 interhomolog recombination mediated by Rad51 when the mechanisms that inhibit Rad51 (*hed1Δ RAD54-T132A*, abbreviated *hedΔR*) are absent (ZIESEL *et al*. 2022). Spore viability of a meiotic depletion allele of *PIF1* (*pif1-md*) is reduced to 80-85% and is not due to MI non-disjunction, similar to *rad51-II3A* (VERNEKAR *et al*. 2021; ZIESEL *et al*. 2022). Taken together these observations suggest that *PIF1* plays a role in Rad51-mediated Phase 2 recombination.

Studying Phase 2 meiotic DSB repair is challenging because the number of DSBs in pachytene cells is low and new DSBs continue to be made (XU *et al*. 1995; ALLERS AND LICHTEN 2001; SUBRAMANIAN *et al*. 2016; MU *et al*. 2020). Therefore, repair of “old” DSBs may be masked by the appearance of new DSBs. To get around this problem, a method using the *NDT80-IN* system was developed to analyze Phase 2 DSB repair by preventing new breaks from occurring during pachynema. Using this assay, as well as co-induction experiments in the *NDT80-IN* system, we have explored the roles of *RAD54*, *TID1* and *PIF1* in Phase 2 DSB repair during yeast meiosis.

## MATERIALS AND METHODS

### Strains and media

All strains were derived from the SK1 background, and their genotypes are listed in Table S1. Growth media are described in (LO AND HOLLINGSWORTH 2011). Sporulation (Spo) medium consisted of 2% potassium acetate.

Gene deletions and *P_GAL1_* fusions were generated by polymerase chain reaction (PCR) based methods using *kanMX6* (pFA6a-*kanMX6*), *natMX4* (p4339) and *hphMX4* (pAG32) as selectable markers that confer resistance to G418, nourseothricin (NAT) and Hygromycin B (HygB), respectively (LONGTINE *et al*. 1998; GOLDSTEIN AND MCCUSKER 1999; TONG *et al*. 2001). All deletions were confirmed by PCR using a forward primer upstream of the open reading frame (ORF) and a reverse primer complementary to the selectable marker. In addition, the absence of the wild-type allele was confirmed using the same forward primer and a reverse primer internal to the ORF. The NH2717 *NDT80-IN pif1-md* diploid was created by first putting the *PIF1* gene under the control of the *GAL1* promoter in the haploid parents of NH2127 as described in (ZIESEL *et al*. 2022). These haploids were then mated to create the diploid. Various *P_GAL1_-PIF1* alleles were introduced in this diploid by digesting either pDM1 or its derivatives with XcmI to target integration at *leu2*. The *LEU2* vector, pRS305, was similarly integrated as a negative control. The presence of the integrated *P_GAL1_-PIF1* alleles was confirmed by PCR using genomic DNA with a forward primer located within the *GAL1* promoter and a reverse primer in the *PIF1* ORF. All of the plasmids were integrated in the diploid and were hemizygous, except for pDM1-m1-K264A-3xFLAG, which was integrated into both haploids that were then mated to make a homozygous diploid as two copies of *pif1-m1-K264A-3xFLAG* are necessary for wild-type protein levels (ZIESEL *et al*. 2022).

The *NDT80-IN REC104-AA* diploid, NH2782, was made by multiple crosses. First, KBY257 was crossed to 7663 to make NH2762. This diploid is heterozygous for *frp1Δ::natMX4*, *SPO11-myc::TRP1*, *RPL13A-2xFKBP12::TRP1 REC104-FRB-3HA-kanMX6* and *yflo22cΔ(1042 to 678)::kanMX4Δ.* NH2762 was sporulated and tetrads dissected. Spore colonies were screened for NAT resistance (*fpr1Δ::natMX4*), Trp^+^ (*SPO11-myc::TRP1* and/or *RPL13A-2xFKBP::TRP1*) and G418 resistance (*REC104-FRB-3HA::kanMX6* and/or *kanMX4::yfl022cΔ(1042to678).* PCR confirmed the presence of *RPL13a-2XFKBP1::TRP1* and *yfl022cΔ(1042to678)::kanMX4* alleles and the absence of *SPO11-myc* in the *MAT***a** haploid, NH2762-24-3. NH2762-24-3 was then crossed to NH2110-7-1 to make NH2763. This diploid is heterozygous for *P_GPD_ - GAL4(848).ER::URA3*, *TRP1-P_GAL1_-NDT80*, *fpr1Δ::natMX4*, *RPL13a-2xFKBP12::TRP1* and *kanMX4::yfl022cΔ(1042 to 678)*. The spore colonies were screened for tetrads that were non-parental ditypes for Trp^+^ (*TRP1-P_GAL1_-NDT80* and *RPL13a-2xFKBP12::TRP1*), Ura^+^ [*P_GPD_-GAL4(848).ER::URA3*], Nat^R^ (*fpr1Δ::natMX4*) and G418^S^ (*YFL022c*). PCR confirmed the presence of *P_GAL1_-NDT80* and *RPL13a-2xFKBP12* in the *MAT***a** haploid, NH2763-39-4. *REC104* was tagged with *FRB-3xHA* by amplifying a 2.7 kb fragment from 7663 that contains 300 bp upstream of the *FRB-3xHA* sequence in *REC104-FRB-3xHA* and 500 bp of 3’ untranslated region (UTR) downstream of *kanMX6*. This fragment was transformed into NH2763-39-4 to make NH2763-39-4 FRB. The presence of the *REC104-FRB-3xHA* fusion was confirmed by PCR. The last step was to introduce the *tor1-1* mutation. NH2763-39-4 FRB was crossed to H6839. Spore colonies of opposite mating type that were Ura^+^ [*P_GPD_ - GAL4(848).ER::URA3*], His^+^ (*tor1-1::HIS3*), Nat^R^ (*fpr1Δ::natMX4*) and G418^R^ (*REC104-FRB-3xHA::kanMX6*) were selected and then screened by PCR for the presence of *TRP1-P_GAL1_-NDT80*. The resulting haploids, NH2781-9-1 and NH2781-22-2 were mated to make NH2782. *REC104* was reintroduced into NH2782 and its derivatives by transforming the diploid with pRD3 (*LEU2 REC104*) digested with SphI to target integration into *REC104-FRB-3xHA*.

Mutant derivatives of NH2782 were made as follows. *RAD54* was substituted with *hphMX4* in NH2781-9-1 and NH2781-22-2 and the resulting haploids mated to make NH2792. Introduction of the *TID1-FRB* and *P_CLB2_-PIF1* (*pif1-md*) alleles required using *kanMX6*. Therefore, an internal deletion of the *kanMX6* gene adjacent to *REC104-FRB-3HA* was created using CRISPR. A healing fragment was generated by annealing complementary primers in which the first 50 bp of the *kanMX6* gene were fused to the last 50 bp. The NH2781-9-1 and NH2781-22-2 haploids were then transformed with the high copy *LEU2* plasmid, pRS425-Cas9-kanMX, which encodes both Cas9 and a guide RNA that targets cutting within the *kanMX6* ORF as well as the healing fragment. Transformants were selected on SD-leu plates. Only those cells in which the DSB was repaired by the healing fragment or by non-homologous end joining were able to grow. As expected, significantly more transformants were observed when the healing fragment was present. Leu^+^ transformants were patched onto YPD plates and screened for G418 sensitivity and loss of the *LEU2* plasmid. The resulting haploids were named NH2781-9-1 Δk and NH2781-22-2 Δk. To introduce the *TID1-FRB* allele, a fragment containing the last 100 bp of *TID1* fused to *FRB::kanMX6* followed by first 100 bp of 3’ UTR was generated by PCR using genomic DNA from H7485. This fragment was transformed into NH2781-9-1 Δk and NH2781-22-2 Δk selecting for G418 resistance. The presence of *TID1-FRB::kanMX6* was confirmed by PCR using genomic DNA from the transformants using the same pair of flanking primers. The two haploids were mated to create NH2814. *RAD54* was deleted from the haploid parents of NH2814 and resulting transformants were mated to make NH2818. *PIF1* was put under the *CLB2* promoter in NH2781-9-1Δk and NH2781-22-2Δk as described above and the haploids mated to make NH2793. The *NDT80-IN REC104-AA rad54Δ TID1-AA pif1-md* diploid NH2839 was made by crossing the *pif1-md* strain, NH2781-9-1 Δk *pif1-md* to a *rad54Δ TID1-AA* strain (NH2822-14-2) (both containing *NDT80-IN* and *REC104-AA*). Haploid segregants of opposite mating type containing *rad54Δ::hphMX4* were selected based on their resistance to HygB, while the presence of the *FRB* tag on *TID1* was confirmed using primers located 100 bp upstream and downstream of the *TID1* stop codon and *pif1-md* was verified using a forward primer in the *kanMX6* gene upstream of the *CLB2* promoter and a reverse primer in the *PIF1* ORF. These haploids were mated to make NH2839. The *NDT80-IN REC104-AA pif1-md rad54Δ* diploid NH2841 was generated by crossing NH2781-9-1 Δk pif1-md to NH2781-22-2 rad54Δ. Haploid segregants of opposite mating type with the *NDT80-IN REC104-AA pif1-md rad54Δ* genotype were mated to make NH2841.

### Plasmids

The genotypes of plasmids used in this work are listed in Table S2. DNA that was amplified by PCR or altered by site-directed mutagenesis was sequenced in the final plasmids by either the Stony Brook University DNA sequencing facility or by Plasmidsauraus (https://www.plasmidsaurus.com/). A *PIF1* inducible allele (*PIF1-IN*) was constructed using the NEBuilder HiFI DNA Assembly Master Mix (hereafter referred to as Gibson Assembly or GA)(New England Biolabs, Cat. # E2621) to fuse the *GAL1* promoter to the *PIF1* gene in the *LEU2* integrating plasmid pRS305 to make pDM1. First a 466 bp fragment containing homology to 25 bp of pRS305 containing the XbaI site fused to the *GAL1* promoter was amplified by PCR using pBG4 as the template. Second, a 2995 bp fragment containing the end of the *GAL1* promoter fused to the *PIF1* open reading frame (ORF), 358 bp of 3’ untranslated region (UTR) and 25 bp of homology including the ApaI site on pRS305 was amplified using pJW5 as the template. The two fragments were assembled into XbaI/ApaI digested pRS305 by GA to make pDM1. To make the *pif1-m1-3xFLAG-IN* and *pif1-m1-K264A-3xFLAG-IN* alleles, the same strategy was used as for pDM1 except that the templates were pJW14 and pJW14-K264A, respectively. The L354P mutation was introduced into *pif1-m1-3xFLAG-IN* by site directed mutagenesis. Complementary 50 bp primers containing CCC (encoding proline) in place of the CUC at codon 354 were used to replicate around each strand of the pDM1-m1-3xFLAG plasmid. The template plasmid was degraded using DpnI and the reaction mixture transformed into *E. coli*. DNA isolated from individual transformants was screened for the presence of the mutation by sequencing. The *REC104 LEU2* integrating plasmid, pRD3, was created first amplifying a 1.3 kb fragment containing the *REC104* ORF with 600 bp 5’ UTR and 140 bp 3’ UTR, as well as overlapping homologies to the ApaI and XbaI sites of pRS305. This fragment was then combined with pRS305 digested with ApaI and XbaI by GA. The CRISPR plasmid, pRS425-Cas9-kanMX contains the *CAS9* gene under the control of the *TEF1* promoter, as well as the sequence 5’TTACTCACCACTGCGATCCC3’ expressed by the *SNR52* promoter to produce a guide RNA that targets cutting ∼300 bp downstream of the *kanMX6* initiation codon.

### Time courses

Cells were streaked for single colonies on YPD plates from freezer stocks incubated at 30°C for 3 days. Single colonies were inoculated into YPD, grown overnight and diluted into YPA in a flask with a minimum YPA:flask volume of 1:5. The cultures were incubated in a 30°C shaker at 240 revolutions per minute (rpm) for several hours. The OD_660_ for 1:1 dilution from each culture was used to determine the cell density. Cells were pelleted and then resuspended in the volume of Spo medium needed for a concentration of 3 x 10^7^ cell/mL. The same amount of sporulating culture was used for all replicates in an experiment. For Southern blots, 10 mL cells were taken at each time point and added to a 50 mL conical tube containing 10 mL 95% ethanol and 1 mL 0.5 M EDTA and stored at -20°C. For 4’,6-Diamidino-2-phenylindole (DAPI) (Vector Laboratories H-1200-10) staining, 0.5 mL cell were fixed with a 1/10 volume of 37% formaldehyde. For protein samples, 5 mL cells at each time point were pelleted and frozen at -80°C. For *NDT80-IN* strains, sporulation was induced by the addition of 5 mM β-Estradiol (Sigma, E8875) to a final concentration of 1 μM. For strains containing *REC104-AA*, a 1 mg/mL stock of Rapamycin (LC Laboratories, R-5000) resuspended in dimethyl sulfoxide (DMSO, Sigma D8418) was added to a final concentration of 1 μg/mL. An equivalent amount of DMSO was added to −Rap cultures as a negative control. A 10 mM stock of the Mek1-as inhibitor, 1-NA-PP1 (Cayman Chemical, 10954) was diluted in sporulating cultures to a final concentration of 1 μM.

### Chromosome spreads

For cytological analysis of Ecm11 localization, 6.7 mL sporulating culture were taken at the indicated timepoints and processed that day. Cells were spheroplasted and spread nuclei were immunostained as described in (GRUBB *et al*. 2015). Surface spread nuclei were incubated with α-Ecm11 antibodies (1:200) at 37°C for 1 hour, followed by incubation with α-guinea pig secondary antibodies containing Alexa-488 at 37°C for 1 hour. Images were collected using a ZEISS Axio Imager.Z2 with a Zeiss Plan-Apochromat 100X objective and analyzed using ZEN 3.5 software.

### Immunoblots and Antibodies

A list of antibodies, their dilutions, and sources is contained in Table S3. Protein extracts were isolated from frozen cell pellets and used for immunoblot analyses as described in (WENG *et al*. 2024). Immunoblot data was collected using a BioRad Chemidoc machine and protein quantification was performed using the Biolab ImageLab software. A custom Ecm11 antibody was generated by Covance Laboratory (now Labcorp) by injection of guinea pigs with the peptide, CDSFDLSRDEKPYIQK, which contains amino acids 128-143 in the 302 amino acid Ecm11 protein.

### Physical analysis of DSBs

Southern blot analysis of DSBs was performed as described in (OWENS *et al*. 2018; GAGLIONE *et al*. 2025).

## RESULTS

### *PIF1* promotes Rad51-mediated repair during yeast meiosis in the absence of DMC1

To test whether *PIF1* can specifically promote Rad51-mediated DSB repair during meiosis, DSB repair was examined in a *dmc1Δ* mutant where Rad51 is the only recombinase available to mediate strand invasion but is prevented from doing so by Hed1 and phosphorylation of Rad54 T132. The resulting unrepaired DSBs trigger the MRC resulting in prophase I arrest (BISHOP *et al*. 1992; TSUBOUCHI AND ROEDER 2006; NIU *et al*. 2009). When these inhibitory mechanisms are removed in a *dmc1Δ hedΔR* diploid, spore viability is ∼70%, demonstrating that Rad51 can generate interhomolog crossovers in the absence of *DMC1* (TSUBOUCHI AND ROEDER 2006; LAO *et al*. 2013; CALLENDER *et al*. 2016). However, spore viability decreases to ∼5% when *PIF1* is meiotically depleted in the *dmc1Δ hedΔR* background, suggesting that *PIF1* promotes Rad51-dependent interhomolog recombination when high levels of Mek1 kinase activity are promoting interhomolog bias (ZIESEL *et al*. 2022).

To determine more directly whether *PIF1* functions in Rad51-mediated DSB repair during meiosis, a *dmc1Δ mek1-as* diploid was used. The *mek1-as* (analog sensitive) allele encodes a kinase with an enlarged ATP pocket that enables inhibition of Mek1-as by addition of the 1-NA-PPI inhibitor to the sporulation (Spo) medium (WAN *et al*. 2004). In the absence of inhibitor, Rad51 activity is inhibited and the cells arrest in prophase I. Inhibition of Mek1-as in a *dmc1Δ mek1-as* strain eliminates Rad54 T132 phosphorylation and results in Hed1 degradation, thereby allowing Rad51-mediated repair of DSBs using sister chromatids and progression through meiosis and sporulation (NIU *et al*. 2005; NIU *et al*. 2009; CALLENDER *et al*. 2016). To determine whether *PIF1* promotes Rad51-mediated DSB repair in the *dmc1Δ* background, the kinetics of DSB disappearance after Mek1-as inhibition was compared between *dmc1Δ mek1-as* and *dmc1 mek1-as pif1-md* strains.

The two diploids were incubated in Spo medium for 5 hours, after which time cells were arrested in prophase I (Figure 1A). When Mek1-as was inhibited, the *dmc1Δ mek1-as* and *dmc1Δ mek1-as pif1-md* cultures progressed through the MI and Meiosis II (MII) with similar kinetics and sporulated to the same extent (Figure 1A, 1B and 1C). In contrast, when DMSO was added to an aliquot of cells from each culture instead of the inhibitor, the cells failed to sporulate as expected (Figure 1C).

**Figure 1.**
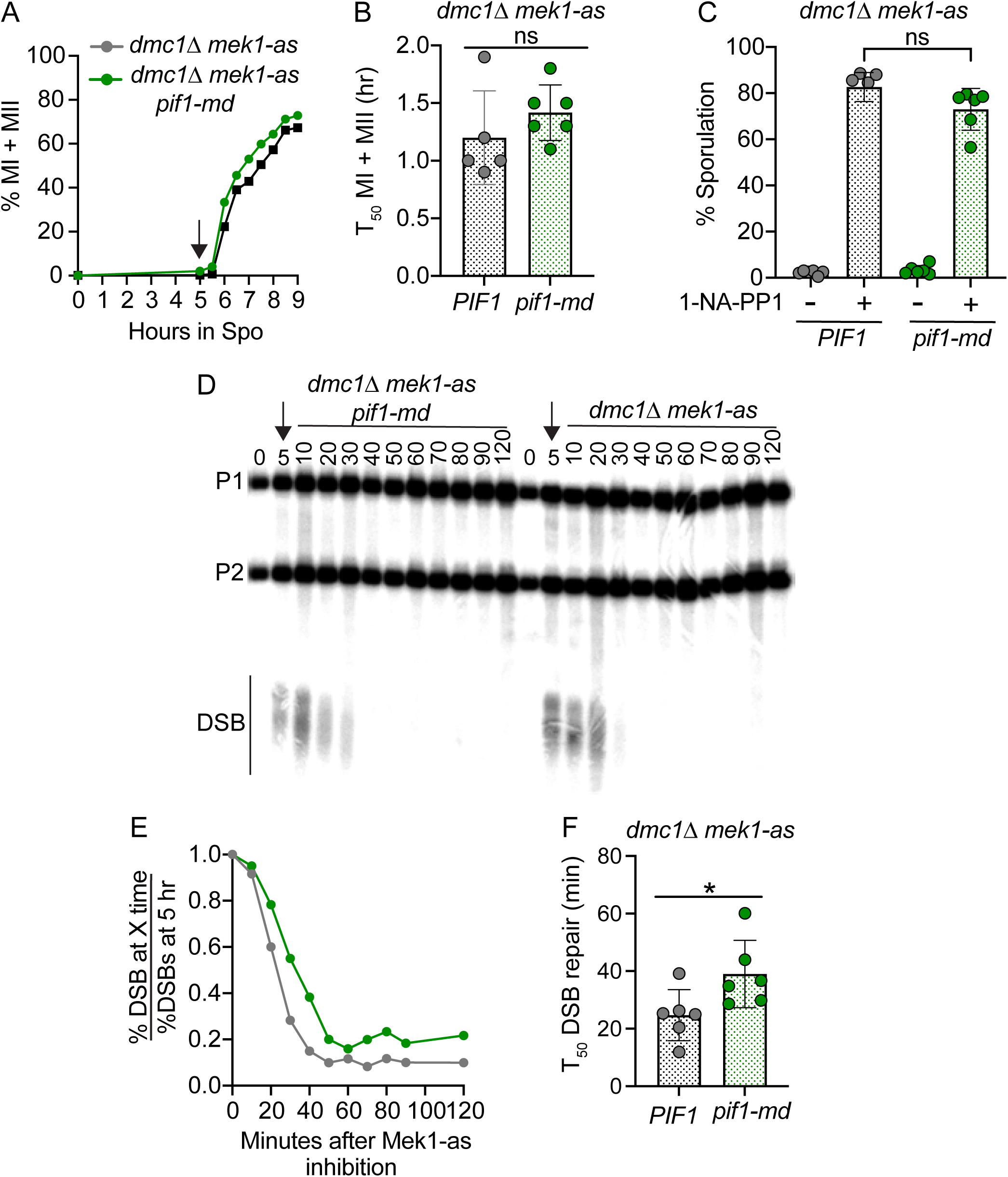
*PIF1* promotes Rad51-mediated intersister DSB repair in a *dmc1Δ* mutant. The *dmc1Δ mek1-as* (NH794::pJR2) and *dmc1Δ mek1-as pif1-md* (NH2754::pJR2) diploids were transferred to Spo medium and incubated for 5 hours at 30°C. A final concentration of 1 μM 1-NA-PP1 or equivalent volume of DMSO was added, after which cells were processed at various timepoints for different phenotypes. (A) Meiotic progression. After addition of inhibitor (indicated by the arrow), cells were taken at 30 minute intervals, fixed and stained with DAPI. For each timepoint, 200 cells were examined by fluorescence microscopy for binucleate and tetranucleate cells that have completed MI and MII, respectively. The average percentage of MI + MII cells was plotted (*n* = 6 for each strain). (B) Meiotic progression timing for each replicate from Panel A was compared using T_50_ values that indicate the amount of time (hr) it took for a sporulation culture to reach one-half the maximum % MI+MII value after addition of 1-NA-PP1. (C) Sporulation of the cultures from Panel A with and without 1-NA-PP1. 200 cells for each replicate were examined by light microscopy for the presence of asci. (D) Representative Southern blot of DNA repair at the *HIS4LEU2* hotspot. Genomic DNA from each timepoint was digested with XhoI to produce two parental fragments (P1 and P2) and DSB fragments created by Spo11. 0 and 5 indicate the hours after transfer to Spo medium, numbers under the line indicate minutes after addition of 1-NA-PP1 (indicated by an arrow). (E) Quantification of the %DSBs at each timepoint was normalized to the %DSBs at 5 hours. The average values for six replicates are plotted. (F) T_50_ analysis of the time in minutes it took to repair one-half the DSBs at 5 hours. The statistical significance of differences between strains in Panels B, C and E was determined using the Mann-Whitney test (*= *p* < 0.05).

DSB repair was monitored by physical analysis of the *HIS4LEU2* hotspot. The DSB sites in this artificial hotspot are flanked by asymmetric XhoI sites. Southern blot analysis of genomic DNA digested with XhoI detected the two parental bands (P1 and P2), as well as smaller fragments resulting from Spo11 DSBs (HUNTER AND KLECKNER 2001). At the 5 hour time point, the arrested cells exhibited hyper-resected DSBs (Figure 1D). After inhibition of Mek1-as, the DSBs took longer to disappear in the *dmc1Δ mek1-as pif1-md* diploid compared to *dmc1Δ mek1-as* (Figure 1E). T_50_ values measured the time it took for half of the DSBs present at 5 hours to disappear. The T_50_ values for *dmc1Δ mek1-as pif1-md* were significantly slower than *dmc1Δ mek1-as*, indicating that *PIF1* promotes Rad51-mediated intersister meiotic DSB repair (Figure 1F).

### *PIF1* promotes Phase 2 recombination in late prophase I

We propose that *PIF1* plays a role in the naturally occurring process of Phase 2 residual DSB repair that occurs during pachynema of prophase I after decreased Mek1 kinase activity results in activation of Rad51. This hypothesis is based on the observation that *pif1-md* spore inviability is similar to that of the strand exchange defective *rad51-II3A* mutant and in both cases the dead spores are not due to MII non-disjunction (Figure 2E and 2F)(CLOUD *et al*. 2012; ZIESEL *et al*. 2022). This idea was tested using an *NDT80* inducible diploid (*NDT80-IN*) (BENJAMIN *et al*. 2003; PRUGAR *et al*. 2017). To look at Phase 2 recombination, *NDT80-IN* diploids were incubated in Spo medium for 5 hours to allow Phase 1 recombination and chromosome synapsis, and *NDT80* transcription was then induced with ED to allow cells to complete Phase 2 recombination, exit prophase I and sporulate. The spore viability of the resulting asci was then determined by tetrad dissection.

**Figure 2.**
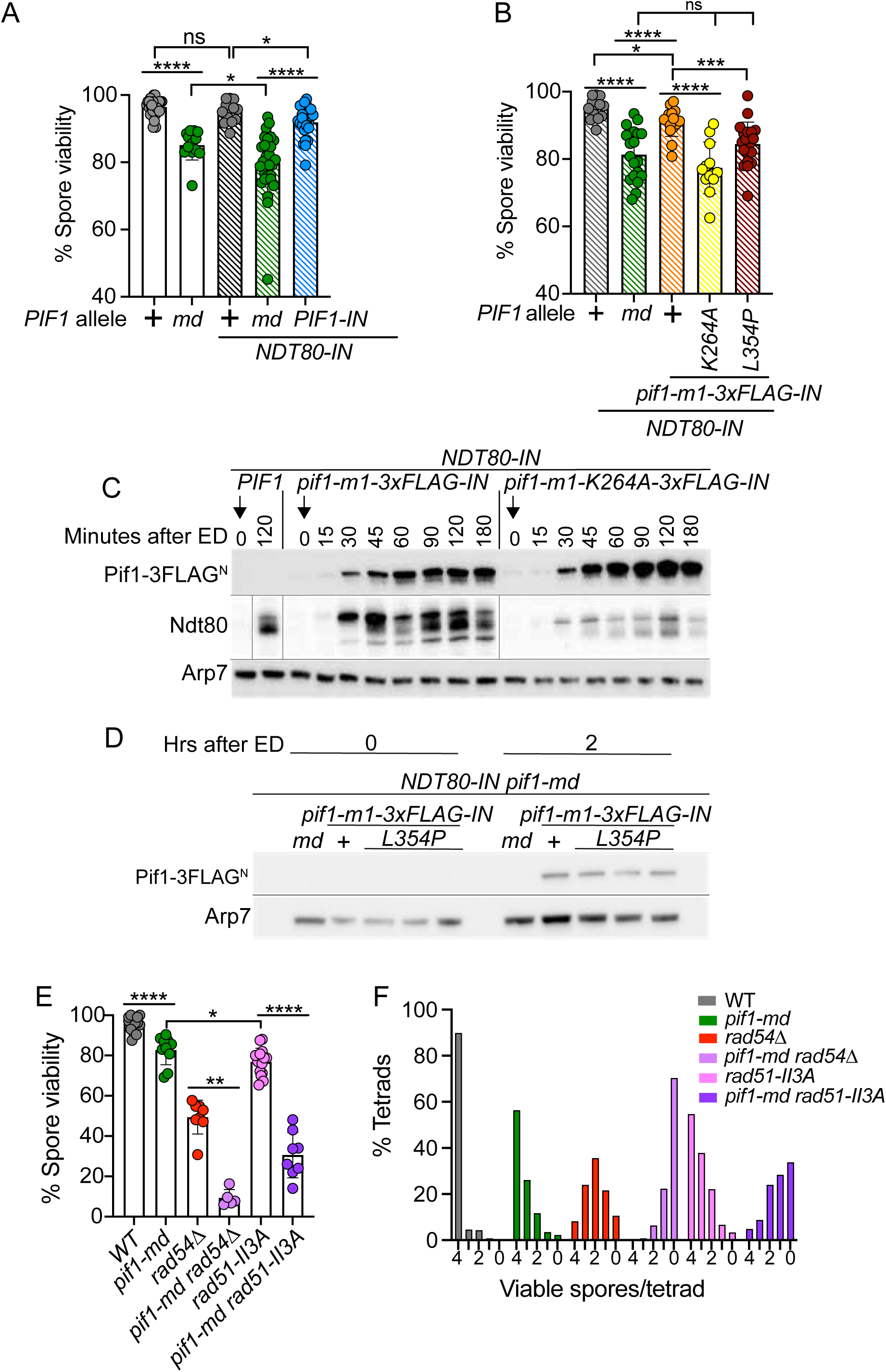
Co-induction of *PIF1* and *NDT80* late in prophase I rescues the *pif1-md* spore inviability phenotype. (A) Co-induction of *NDT80* and *PIF1*. Diploids containing *NDT80-IN* (NH2127, *n* = 3), *NDT80-IN pif1-md* (NH2717, *n* = 4) or *NDT80-IN pif1-md::PIF1-IN* (NH2717::pDM1, *n* = 3) were incubated in Spo medium for 5 hours at which time a final concentration of 1 μM ED was added to induce *NDT80* and *PIF1* and allow cells to sporulate. Spore viability was determined by tetrad dissection. Each dot represents a plate containing at least 20 tetrads combining technical replicates of each biological replicate (*n*). “+” indicates a wild-type allele. Hatch marks indicate *NDT80-IN* diploids. Data for the wild-type and *pif1-md* diploids were taken from (ZIESEL *et al*. 2022). (B) Co-induction of *NDT80* and various *pif1* alleles*. NDT80-IN pif1-md* (NH2717::pRS305, *n* = 2), *NDT80-IN pif1-m1-3xFLAG* (NH2717:: pDM1-m1-3xFLAG, *n* = 3), *NDT80-IN pif1-m1-K264A-3xFLAG* (NH2717:: pDM1-m1-K264A-3xFLAG^2^, *n* = 3) and *NDT80-IN pif1-m1-L354P-3xFLAG* (NH2717:: pDM1-m1-L354P-3xFLAG, *n* = 3) were sporulated and dissected as in Panel A. The *NDT80-IN* data are the same as Panel A. (C) Kinetics of Ndt80, Pif1-3xFLAG and Pif1-K264A-3xFLAG protein induction. *NDT80-IN*, *NDT80-IN pif1-m1-3xFLAG,* and *NDT80-IN pif1-m1-K264A-3xFLAG* were incubated in Spo medium for 5 hours at which time ED was added (indicated by arrows). Protein extracts from cells taken at 15 minute intervals after addition of ED were probed with α-Ndt80 and α-FLAG antibodies to detect Ndt80 and Pif1, respectively. Arp7 was used as a loading control. Gray lines indicate places where irrelevant lanes were removed or different blots were used. This experiment was repeated twice with similar results. (D) Comparison of Pif1-3xFLAG^N^ and Pif1-L354P-3xFLAG^N^ steady state protein levels before and after addition of ED after 5 hours in Spo medium. One replicate from *NDT80-IN pif1-md* (*md*) and *NDT80-IN pif1-m1-3xFLAG* (indicated by “+”) are shown as well as three biological replicates for *NDT80-IN pif1-m1-L354P-3xFLAG*. (E) Epistasis analysis. Single colonies from wild type (NH716), *pif1-md* (NH2657), *rad51-II3A* (NH2618), *rad54Δ* (NH1018), *pif1-md rad51-II3A* (NH2743 RCEN) and *pif1-md rad54Δ* (NH2785) were sporulated on solid medium and spore viability determined as in Panel A. Statistical differences between strains shown in Panels A and B were determined using the Mann-Whitney test (* = *p* < 0.5; ** = *p* < 0.1; *** = *p* < 0.001, **** = *p* < 0.0001). (F) The distribution of viable spores in tetrads for the dissections shown in Panel E. The total number of tetrads is wild type (388), *pif1-md* (257), *rad54Δ* (208), *pif1-md rad54Δ* (125), *rad51-II3A* (372), and *pif1-md rad51-II3A* (204).

In the *NDT80-IN pif1-md* diploid, Pif1 was absent during both Phase 1 and Phase 2 and spore viability was decreased to a level similar to *pif1-md* (Figure 2A). Like *pif1-md*, the distribution of viable spores in tetrads in the *NDT80-IN pif1-md* diploid did not exhibit an MI non-disjunction pattern (a decrease in tetrads with four viable spores with a corresponding increase in 2 and 0 viable spore tetrads)(Figure S1A) (HOLLINGSWORTH *et al*. 1995; ZIESEL *et al*. 2022). To determine if *PIF1* is specifically required for Phase 2 recombination, *P_GAL1_-PIF1* was integrated into the *NDT80-IN pif1-md* diploid so that transcription of both *NDT80* and *PIF1* were induced at the same time by ED. For simplicity, the genotype of this diploid is referred to as *NDT80-IN PIF1-IN*. Co-induction of *PIF1* and *NDT80* significantly increased spore viability to almost wild-type levels by raising the number tetrads in which all four spores were viable (Figures 2A and S1A). These results suggest that *PIF1* has a role in Phase 2 recombination to repair residual DSBs.

The interpretation of the *PIF1*/*NDT80* co-induction experiment assumes that the two proteins were generated with the same kinetics. This assumption was tested using a modified *P_GAL1_-PIF1* allele that encodes three FLAG epitopes at the C-terminus of the helicase to allow detection of the protein on immunoblots. Pif1 is present in two isoforms in the cell, one is nuclear, while the other protein isoform is targeted to mitochondria by an N-terminal signal sequence (IVESSA *et al*. 2000). Mutation of the initiating methionine removes the signal sequence resulting in predominantly nuclear Pif1 (*P_GAL1_-pif1-m1-3xFLAG*, which is referred to *pif1-m1-3xFLAG-IN* (SCHULZ AND ZAKIAN 1994). For simplicity, the protein encoded by this allele is referred to as Pif1-3xFLAG^N^ (N = nucleus). The *pif1-m1-3xFLAG-IN* allele was integrated into the *NDT80-IN pif1-md* diploid where it complemented the spore viability defect of the *NDT80-IN pif1-md* mutant to nearly wild-type levels (Figure 2B). (All plasmid borne *PIF1* alleles were integrated into *pif1-md* strains. For simplicity, the genotypes refer only to the allele on the plasmid.)

The expression kinetics of the Pif1-3xFLAG^N^ and Ndt80 proteins was analyzed in protein extracts taken from an *NDT80-IN pif1-m1-3xFLAG-IN* strain at various times after addition of ED. The *NDT80-IN* control exhibited Ndt80 protein 2 hours after induction, but no Pif1 as expected (Figure 2C). In the *NDT80-IN PIF1-IN* diploid, both Ndt80 and Pif1-3xFLAG^N^ first appeared weakly 15 minutes after induction and protein levels peaked by 90 minutes. Importantly, there was no evidence that Pif1-3xFLAG^N^ was present earlier than Ndt80.

Pif1 helicase activity is required both for break-induced replication in vegetative cells and for the promotion of Rad51-mediated interhomolog recombination during Phase 1 recombination of meiosis (WILSON *et al*. 2013; ZIESEL *et al*. 2022). The *pif1-K264A* mutation eliminates helicase activity (SCHULZ AND ZAKIAN 1994). The *pif1-m1-K264A-IN* mutant failed to complement the spore inviability of *NDT80-IN pif1-md* (Figures 2B and S1B). Steady state Pif1-K264A-3xFLAG^N^ protein levels were similar to Pif1-3xFLAG^N^ and similar kinetics was observed between Ndt80 and Pif1-K264A-3xFLAG^N^, although the amount of Ndt80 was reduced in this strain compared to the *NDT80-IN pif1-m1-3FLAG-IN* diploid for unknown reasons (Figure 2C).

In mammalian tissue culture cells, break-induced replication is defective in *a pif1* mutant that alters the 319th amino acid from leucine to proline and this mutation is associated with breast cancer in humans (CHISHOLM *et al*. 2012). Importantly, L319 is located within the Pif1 family specific sequence (PFSS) present in an α-helix that is important for holding a pin-loop structure in place that directs the unwinding of the double helix (Figure S1C) (CHEN *et al*. 2016; LU *et al*. 2018). L319 in human Pif1 corresponds to L354 in yeast Pif1 (Figure S1D).

The spore viability of an *NDT80-IN pif1-m1-L354P-3xFLAG-IN* diploid was decreased to a similar level as *NDT80-IN pif1-md*, indicating a failure to complement (Figures 2B and S1B). Importantly, steady state levels of the Pif1-L354P-3xFLAG^N^ protein were not decreased compared to Pif1, ruling out a trivial explanation for the mutant phenotype (Figure 2D). Therefore, similar to break-induced replication, Pif1 helicase activity is required for Phase 2 meiotic recombination in yeast.

### *PIF1* function in promoting Phase 2 recombination is not dependent upon *RAD51* or *RAD54*

All our results thus far are consistent with a simple model in which Rad51 functions with Rad54 to make D-loops that are then extended by Pif1 to promote the repair of residual DSBs during Phase 2. This model predicts that both *rad51-II3A* and *rad54Δ* should be epistatic to *pif1-md* with regard to spore viability. That is, if no D-loops are generated by Rad51, then *PIF1* should be irrelevant. This idea was tested by assessing the spore viability of *pif1-md rad51-II3A* and *pif1-md rad54Δ* diploids. As has been previously observed, the *rad54Δ* spore viability was significantly worse than the *rad51-II3A* mutant, indicating that *RAD54* function is not limited to Rad51 (SHINOHARA *et al*. 1997; CLOUD *et al*. 2012). In contrast to our expectation, *pif1-md* exhibited a synergistic reduction in spore viability with both *rad54Δ* and *rad51-II3A* that was not due to MI non-disjunction (Figure 2E and 2F). These results suggest that *RAD51*/*RAD54*-independent recombination can occur in Phase 2 that is facilitated by *PIF1*.

### A system for studying Phase 2 meiotic DSB repair

During pachynema, Phase 2 repair of “old” DSBs is offset by the introduction of new DSBs (Figure 3A). Therefore, to study Phase 2 recombination, it is necessary to first turn off DSB formation in *NDT80-IN* pachytene-arrested cells and then allow time for DSB repair to occur. If the repair is efficient, little to no old DSBs should be present when the cells are induced to sporulate by the addition of ED (Figure 3A). The *NDT80-IN* strain was therefore modified to conditionally deplete Rec104, a subunit of the Spo11 core complex, from the nuclei of *ndt80*-arrested cells using the “anchor away” approach to prevent new DSB formation (HARUKI *et al*. 2008; CLAEYS BOUUAERT *et al*. 2021; PRIELER *et al*. 2021). Addition of rapamycin (Rap) results in the specific removal of Rec104-FRB from the nucleus. (For simplicity, *REC104-AA* will be used to indicate that a strain contains *REC104-FRB*, as well as the other genes necessary for the depletion). Cells are then kept in pachynema for an additional two hours to allow DSB repair to occur. The presence of DSBs at nine hours can then be monitored indirectly by Hop1 phosphorylation (BISHOP 1994; CARBALLO *et al*. 2008; CHUANG *et al*. 2012). In addition, cells can be induced to sporulate by addition of ED, thereby allowing indirect detection of DSBs by the highly sensitive spore viability assay.

**Figure 3.**
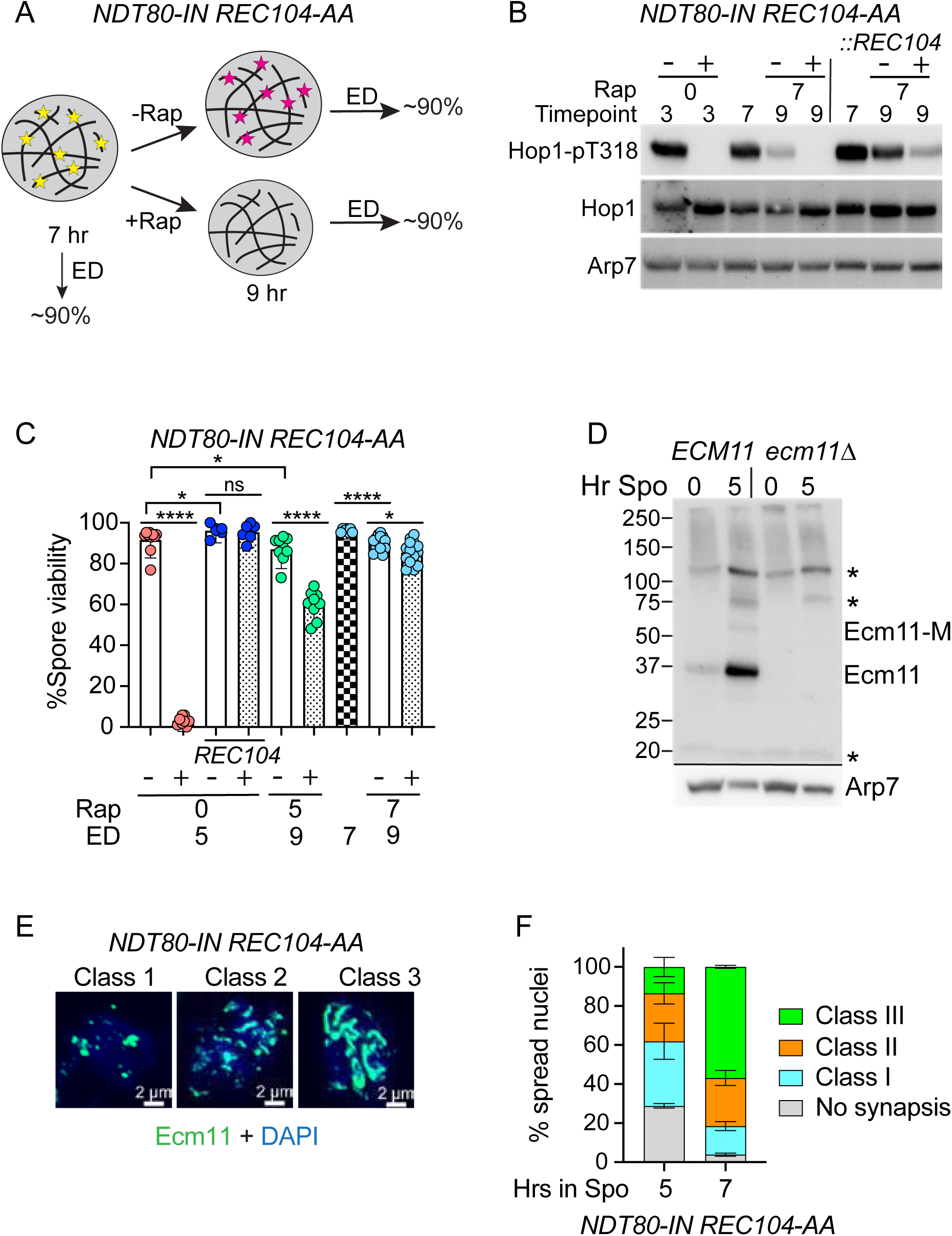
A system for studying the Phase 2 repair of residual meiotic DSBs. (A) Schematic of DSB behavior in pachytene-arrested cells with and without *REC104*. Left side shows an *NDT80-IN REC104-AA* diploid cell arrested in pachynema after 7 hours in Spo medium with fully synapsed homologs and a low number of DSBs (yellow stars= “old” DSBs). Addition of ED results repair of the old DSBs, exit from prophase I and sporulation. Numbers indicate predicted spore viabilities. Right, in the absence of rapamycin (−Rap), after 9 hours Rec104 is present and a low level of new DSBs have been formed (magenta stars), while the old breaks have been repaired in the two hour window. Addition of ED results in repair of the new DSBs, exit from prophase I and sporulation. Addition of Rap (+Rap) at 7 hours depletes Rec104 from the nucleus preventing new DSB formation while repair of the old DSBs still occurs. Addition of ED results in repair of any remaining DSBs, exit from prophase I and sporulation. (B) Indirect detection of DSBs using an antibody specific for Hop1-pT318. To deplete Rec104 from the nucleus, a final concentration of 1 μg/mL Rap was added to the *NDT80-IN REC104-AA* diploid, NH2782, as well as the same diploid containing an integrated copy of *REC104* (NH2782::pRD3) after 0 or 7 hours in Spo medium. “+” indicates Rap, “-“ indicates the addition of an equivalent volume of DMSO and the number under the line indicates the time at which both were added. Protein extracts were made from cells taken at either 3, 7 or 9 hours in Spo. Immunoblots were probed with α-Hop1-pT318, α-Hop1 or α-Arp7. (C) Spore viability of *NDT80-IN REC104-AA* or *NDT80-IN REC104-AA::REC104* with and without addition of Rap at 0, 5 or 7 hours. The speckled pattern indicates the presence of Rap. The checkerboard pattern indicates that neither DMSO nor Rap were added. Cells were induced to sporulate by addition of ED at either 5, 7 or 9 hours. Spore viability was analyzed as in Figure 2. (D) Validation of a new Ecm11 antibody. *ECM11* (NH2436) and *ecm11Δ* (NH2406) diploids were transferred to Spo medium and protein extracts generated from cells at 0 and 5 hours. Immunoblots of these extracts were probed with either α-Ecm11 or α-Arp7 antibodies. Numbers indicate the positions of the prestained molecular weight markers. Asterisks indicate non-specific bands also present in the *ecm11Δ*. Ecm11-M indicates an Ecm11 protein with slower mobility. The black lines show that two different blots were used to probe the extracts for Ecm11 and Arp7. (E) Different stages of chromosome synapsis determined by Ecm11 staining. Representative images show three different classes of Ecm11 staining. Class 1 contains dots indicative of leptotene. Class 2 contains dots and lines indicative of zygotene. Class 3 contains primarily lines indicative of pachytene. (F) Quantification of the degree of synapsis in the wild-type diploid after 5 or 7 hours in Spo medium. For each condition, 130 chromosome spreads from two biological replicates were stained with Ecm11 and classified as either having no synapsis, Class 1, Class 2 or Class 3. Error bars indicate the range.

To test whether addition of Rap to the *NDT80-IN REC104-AA* diploid eliminated DSB formation as expected, an *NDT80-IN REC104*-AA culture was split in half immediately after transfer to Spo medium and DMSO (referred to throughout as −Rap) was added to one culture while Rap (+Rap) was added to the other. Protein extracts were made from cells three hours later and probed with antibodies specific for phosphorylated Hop1-T318 (pT318)(CHUANG *et al*. 2012). Antibodies that detect total Hop1 and Arp7 were used as loading controls. A robust Hop1-pT318 band was observed at 3 hours in the absence, but not the presence, of Rap (Figure 3B). These cultures were induced to sporulate by addition of ED at 5 hours and the resulting tetrads dissected. Spore viability was <5% in the cells treated with Rap, in contrast to the high spore viability of the −Rap cells (Figure 3C). Adding Rap at 0 hours had no effect on spore viability when *REC104* was integrated into the *NDT80-IN REC104-AA* strain (Figure 3C). Taken together, these results demonstrate that Rec104-FRB was sufficiently depleted from meiotic nuclei to prevent DSB formation. Note however, that in the absence of Rap, the *NDT80-IN REC104-AA* diploid exhibited a slightly lower, but statistically significant, decrease in spore viability compared to *NDT80-IN REC104-AA::REC104*, suggesting that the *FRB* tag on *REC104* slightly lowers its functionality (Figure 3C).

Phase 2 recombination occurs after the bulk of DSBs have been processed into double Holliday junctions or non-crossovers and chromosomes have synapsed so additional DSBs are no longer necessary for homolog pairing or making crossover precursor intermediates. Depleting Rec104 from pachytene cells should therefore have no effect on chromosome segregation or spore viability. It is critical, however, that cells have reached pachynema prior to Rec104-FRB removal, otherwise the decrease in DSBs in cells still in leptonema or zygonema would reduce the number of crossovers in those cells, resulting in lower spore viability. Initially, *NDT80-IN REC104-AA* cells were incubated in Spo medium for 5 hours at which time Rap was added to Anchor Away Rec104-FRB. ED was then added at 9 hours to induce sporulation. The spore viability of the −Rap control induced with ED at 9 hours was only slightly lower than the −Rap control to which ED was added at 5 hours (Figure 3C). In contrast, spore viability was significantly reduced when Rec104-FRB was depleted at 5 hours and then induced to sporulate at 9 hours, indicating that many cells had not reached the pachytene stage by 5 hours (Figure 3C).

To determine the fraction of cells in pachytene, chromosome spreads from the 5 hour timepoint of the *NDT80-IN REC104-AA* culture were stained with a newly generated custom antibody that recognizes Ecm11, a meiosis-specific protein component of the central element of the SC (HUMPHRYES *et al*. 2013). This antibody was validated by probing immunoblots of protein extracts from *ECM11* and *ecm11Δ* diploids 0 and 5 hours after transfer to Spo medium. The *ECM11* diploid exhibited a meiotically-induced band of ∼37 kD, in good agreement with Ecm11’s predicted molecular weight of 34 kD (Figure 3D). In addition, an Ecm11-specific band of slower mobility was observed (Ecm11-M), consistent with a previously observed sumoylated form of Ecm11 (Figure 3D)(HUMPHRYES *et al*. 2013). Importantly, these bands were absent in the *ecm11Δ*, confirming the specificity of the antibody (Figure 3D).

The *NDT80-IN REC104-AA* chromosome spreads exhibiting Ecm11 staining were separated into 3 classes, Class 1 exhibits foci (leptonema), Class 2, foci and lines (zygonema) and Class 3, lines (pachynema) (Figure 3E). Sixty percent of the spreads from the 5 hour timepoint exhibited either no synapsis or Class 1 foci, showing that nuclear depletion of Rec104-FRB at this time was premature (Figure 3F). In contrast, when cells were incubated for 7 hours, >80% of the spreads were either Class 2 or Class 3 (Figure 3F). Depletion of Rec104-FRB at 7 hours, followed by ED induction of *NTD80* at 9 hours, resulted in spore viability that was nearly the same as the −Rap control (Figure 3C). Therefore, for all further experiments, cells were incubated in Spo medium for 7 hours, at which time Rap was added to deplete Rec104-FRB, followed by 2 hours to allow Phase 2 DSB repair and then addition of ED at 9 hours to induce sporulation. All strains in the remaining experiments contain *NDT80-IN REC104-IN*. For simplicity, this genotype is inferred, e.g. *NDT80-IN REC104-IN rad54Δ* is referred to as *rad54Δ*.

### New DSB formation masks the repair of DSBs at pachynema

Hop1-pT318 phosphorylation was observed in the wild-type diploid after 7 hours in Spo, confirming the presence of DSBs in pachytene arrested cells (Figure 3B). Inducing *NDT80* at this time results in disassembly of the SC, crossover formation and repair of residual DSBs, resulting in high spore viability (Figure 3C) (SOURIRAJAN AND LICHTEN 2008; PRUGAR *et al*. 2017). Previous work has shown that when Rad54 is anchored away in pachytene arrested cells, the number of Rad51 foci increases (SUBRAMANIAN *et al*. 2016). This increase is dependent upon Spo11 generated breaks, suggesting that DSB repair can occur during pachynema prior to *NDT80* induction. The idea that the DSBs detected at 9 hours were formed after 7 hours is supported by the observation that these breaks were gone when Rec104 was depleted at 7 hours (Figure 3A and 3B). In the diploid containing the integrated *REC104* allele, there are three functional copies of *REC104*, and the Hop1-pT318 signal was stronger at 9 hours in the −Rap condition than that of the *REC104-A*A homozygous diploid (Figure 3B). Importantly when Rec104-FRB was depleted from this diploid at 7 hours, Hop1-pT318 was still detected at 9 hours albeit at a lower level, due to continued DSB formation by Spo11. This experiment shows that at the *ndt80* arrest the number of DSBs present at any given time is the result of a balance between DSB formation and repair. When Rec104 was depleted at 7 hours to eliminate the creation of new DSBs, the old DSBs present at that time were repaired by 9 hours, hence the absence of Hop1 phosphorylation. This method can therefore be used to identify mutants defective in Phase 2 meiotic recombination by detection of DSBs at the 9 hour timepoint after Rec104 depletion at 7 hours.

### Phase 2 meiotic DSB repair can occur independently of *RAD54*

Co-induction of *RAD54* and *NDT80* suppresses the spore inviability of a *rad54Δ*, consistent with the idea that *RAD54* functions specifically in Phase 2 recombination (Figure 2)(PRUGAR *et al*. 2017). To determine whether *RAD54* is required for Phase 2 DSB repair, a *rad54Δ* diploid was examined using our Phase 2 recombination protocol.

An *NDT80 REC104 rad54Δ* diploid produces 50-60% viable spores (SHINOHARA *et al*. 1997). A similar decrease in spore viability was observed when the *rad54Δ* −Rap diploid was induced to sporulate with ED after 9 hours (Figure 4C). The presence of Hop1-pT318 confirmed the presence of DSBs after 7 hours in Spo in this strain (Figure 4B). If all Phase 2 DSB repair requires *RAD54*, then in the −Rap cells, DSBs should accumulate at the 9 hour timepoint, as new breaks are formed, while the old breaks remain. In this case spore viability should go down in cells induced to sporulate at 9 hours (Figure 4A, left). In the +Rap cells, the lack of Rec104 would prevent new DSB break formation, while the old DSBs persist at 9 hours, leading to a similar level of viable spores as those cells induced to sporulate at 7 hours (Figure 4A, left). Alternatively, if some Phase 2 DSB repair can occur independently of *RAD54*, then by 9 hours in the −Rap cells, the DSBs present at 7 hours will have been repaired, and only new DSBs will be detected. How these new breaks would affect spore viability depends upon how frequently DSBs are made and repaired, making the expected number difficult to predict. However, after Rec104 depletion at 7 hours, if DSBs can be fixed without *RAD54*, Hop1-pT318 phosphorylation should decrease at the 9 hour timepoint as the old breaks are repaired and no new DSBs are made (Figure 4A, right). As a result, spore viability should improve.

**FIGURE 4.**
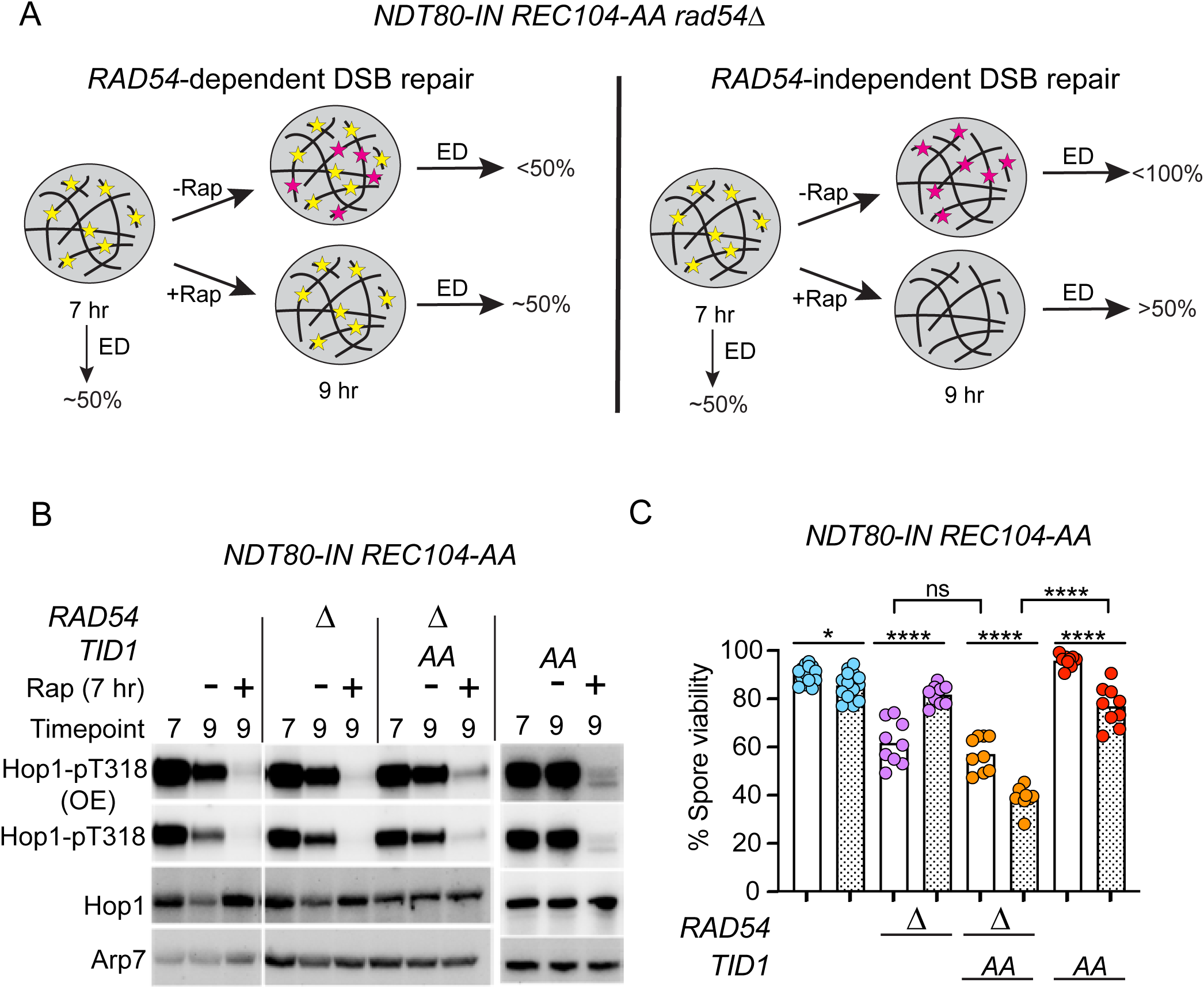
*TID1* promotes Phase 2 DSB repair in the absence of *RAD54*. (A) Schematic showing the expected results based on two alternative hypotheses. Yellow stars indicate “old” DSBs present in pachytene-arrested cells after 7 hours in Spo medium. Red stars indicate new DSBs formed between 7 hours and 9 hours in the absence of presence of Rap. The predicted spore viabilities are shown after inducing sporulation with ED at the indicated times. Left: DSB repair is completely dependent upon *RAD54*. Right: *RAD54*-independent DSB can occur. (B) Indirect detection of DSBs in wild type, *rad54Δ* (NH2792), *rad54Δ TID1-AA* (NH2818) or *TID1-AA* (NH2814) diploid using phosphorylation of Hop1-318 (Hop1-pT318). Protein extracts from the indicated times were probed on immunoblots with antibodies against total Hop1, Hop1-pT318 or Arp7. (C) *NDT80-IN REC104-AA* diploids with the indicated genotypes were incubated in Spo medium for 7 hours at which time either DMSO or RAP (stippled bars) was added. After 2 more hours, sporulation was induced with ED and the resulting tetrads dissected and analyzed as in Figure 2A. “Δ” = *rad54Δ*, “*AA*” indicates *TID1-AA*.

The Hop1 phosphorylation and spore viability assays both indicated that most DSBs were repaired during Phase 2 without *RAD54*. Hop1 phosphorylation was greatly reduced at 9 hours in cells treated with Rap compared to the −Rap control (Figures 4B and S2). Furthermore, spore viability in the +Rap *rad54Δ* cells increased by over 20% compared to the −Rap cells (Figure 4C).

Otherwise wild-type diploids deleted for both *RAD54* and its paralog, *TID1*, exhibit synergistic defects in interhomolog recombination and spore viability, suggesting that *RAD54* can partially substitute for *TID1* during Phase 1 recombination in the absence of the paralog (SHINOHARA *et al*. 1997). We tested whether the reverse is true—that is, can *TID1* promote Phase 2 DSB repair in the absence of *RAD54*? Because *TID1* is involved in Phase 1 repair, a *TID1-FRB* allele was introduced into the *NDT80-IN REC104-AA* background so that the Tid1-FRB protein would be co-depleted with Rec104-AA from pachytene-arrested cells after Phase 1 recombination. There was a slight reduction in spore viability in the *TID1-AA* +Rap cells after addition of Rap at 7 hours, but very little Hop1 phosphorylation, consistent with the idea that Rad54 is normally responsible for the bulk of Phase 2 DSB repair (Figures 4C and S2). The spore viability of the *rad54Δ TID1-AA* diploid in the absence of Rap was similar to *rad54Δ*, as expected since Tid1-FRB is present in the nucleus. However, in the presence of Rap, *rad54Δ TID1-AA* spore viability was reduced and Hop1-pT318 phosphorylation was increased (Figures 4B, 4C and S2). These results demonstrate that *TID1* can promote Phase 2 DSB repair when *RAD54* is absent.

### *PIF1* is required for Phase 2 DSB repair in the absence of *RAD54* and *TID1*

Epistasis analysis between *pif1-md* and both *rad51-II3A* and *rad54Δ* revealed that the function of *PIF1* in Phase 2 recombination is more complicated than our original hypothesis that Pif1 promoted extension of the invading strand from a Rad51-mediated D-loop (Figure 2E). We therefore examined *pif1-md* alone and in combination with *rad54Δ* or *rad54Δ TID1-AA* using our Phase 2 recombination method. The *pif1-md* −Rap diploid showed a small, but significant decrease in spore viability compared to the −Rap wild-type strain, similar to what was previously observed in *pif1-md NDT80 REC104* and *pif1-md NDT80-IN* strains (Figures 2A, 2B and 5C)(ZIESEL *et al*. 2022). Preventing new DSB formation by the addition of Rap had no effect on sporulation or spore viability (Figure 5A and 5B). One explanation is that when Pif1 is depleted during meiosis, a low level of DNA damage occurs during premeiotic S phase that can persist into prophase I and be rescued by induction of *PIF1* during pachynema (Figures 2A and S1A). Alternatively, *PIF1* may function directly in Phase 2 recombination.

**Figure 5.**
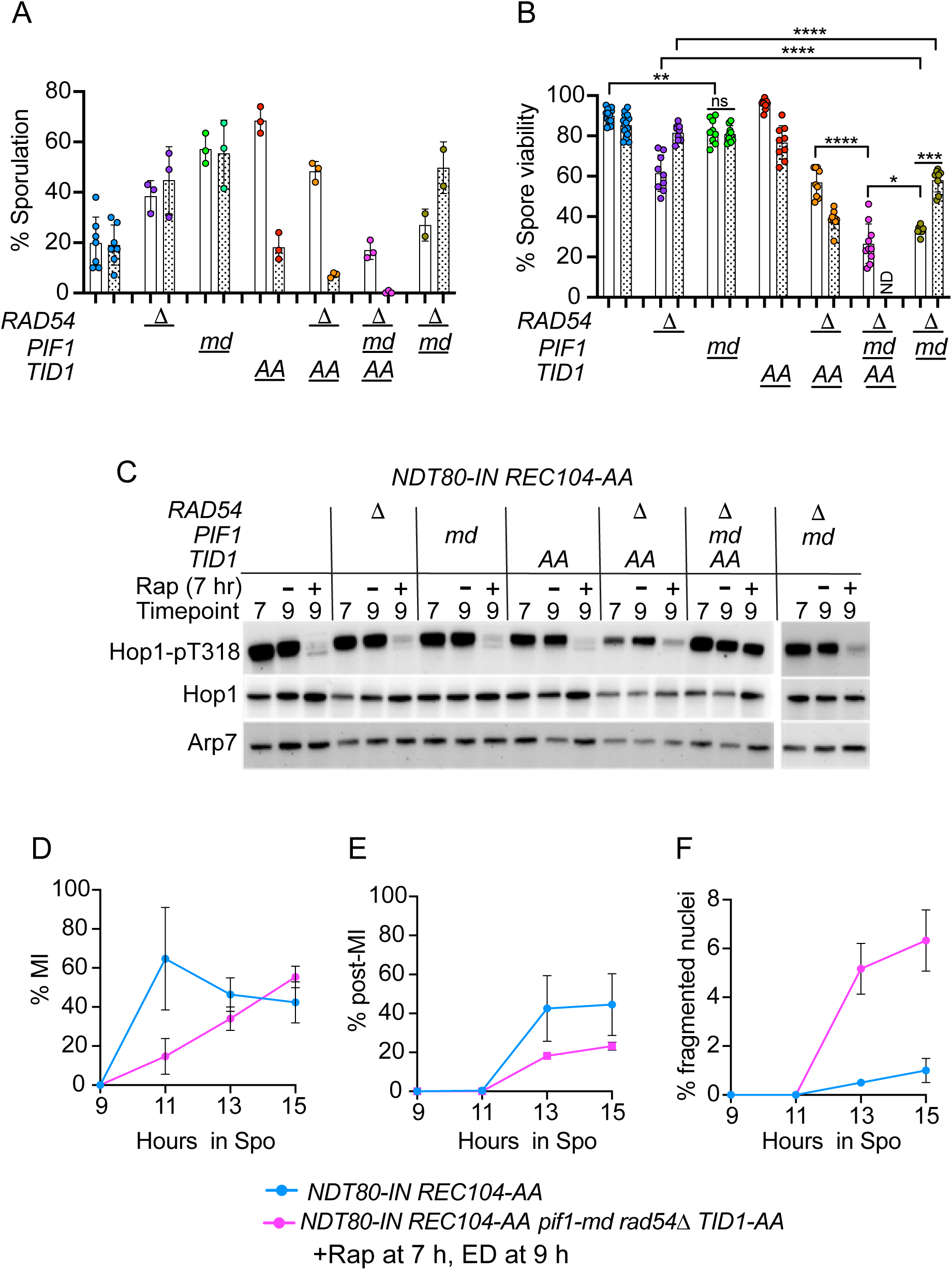
Phase 2 recombination in strains with various combinations of *rad54Δ*, *TID1-AA* and *pif1-md*. All diploids were incubated in Spo medium for 7 hours, DMSO or Rap was added (indicated by stippled bars), and incubated for 2 more hours before inducing sporulation by addition of ED. (A) Sporulation. Wild-type diploids wild-type (NH2782, *n*=7), *rad54Δ* (NH2792, n=3), *pif1-md* (NH2793, *n*=3), *TID1-AA* (NH2814, *n*=3), *rad54Δ TID1-AA* (NH2818, *n*=3), *pif1-md rad54Δ TID1-AA* (NH2839, *n*=3) or *pif1-md rad54Δ* (NH2841, *n*=2) were analyzed for sporulation. Each dot represents a different biological replicate for which 200 cells were counted for the percentage of asci using light microscopy. Error bars indicate the mean and standard deviation, except for *pif1-md rad54Δ* where the range is shown. (B) Spore viability from the cultures used for Panel A was analyzed as described in Figure 2A. “ND” means no data since the cells did not form asci. For comparison, the data for the wild-type, *rad54Δ*, *TID1-AA* and *rad54Δ TID1-AA* strains are repeated from Figure 4C. (C) Hop1-pT318 phosphorylation was analyzed for the seven strains as described in Figure 4C. Different biological replicates are shown for the wild-type, *rad54Δ*, *TID1-AA* and *rad54Δ TID1-AA* strains compared to Figure 4B. (D) Meiotic progression in wild type or *pif1-md rad54Δ TID1-AA* diploids treated with Rap at 7 hours and ED at 9 hours. Cells were stained with DAPI and the average % MI, post MI (includes cells with 4 and >4 DAPI bodies), and % fragmented nuclei (% >4 DAPI bodies) for three biological replicates for 200 cells. Statistical differences between strains shown in Panels A, B, D and E and F were determined using the Mann-Whitney test (* = *p* < 0.5; ** = *p* < 0.1; *** = *p* < 0.001, **** = *p* < 0.0001).

The *pif1-md rad54Δ* −Rap diploid exhibited reduced spore viability below that of either the *rad54Δ* or *pif1-md* −Rap strains, consistent with the epistasis results (Figures 2E and 5B). Both sporulation and spore viability were improved in the *pif1-md rad54Δ* +Rap diploid relative to the −Rap condition, similar to *rad54Δ* alone (Figures 4C, 5A and 5C). Therefore, *PIF1* is not essential for *RAD54*-independent Phase 2 repair. However, the rescue in the +Rap condition was not complete, as the +Rap *pif1-md rad54Δ* diploid exhibited fewer viable spores and more Hop1 phosphorylation than *rad54Δ* +Rap, revealing that some unrepaired DSBs remained in the *pif1-md rad54Δ* +Rap diploid at 9 hours (Figures 5B, 5C and S2).

While depleting *PIF1* did not prevent Phase 2 DSB repair in the *rad54Δ*, addition of *pif1-md* to the *rad54Δ TID1-AA* strain had a drastic effect. In the absence of Rap, Tid1-FRB was still present in the nucleus so the −Rap *pif1-md rad54Δ TID1-AA* diploid should resemble the *pif1-md rad54Δ* strain. In fact, sporulation and spore viability were only slightly reduced between the two −Rap diploids (Figure 5A and 5B.) However, in the +Rap condition, sporulation was abolished in the *pif1-md rad54Δ TID1-AA* diploid so no spore viability data could be obtained (Figure 5A). In addition, Hop1-pT318 phosphorylation was greatly increased, indicating the persistence of a high number of DSBs at the 9 hour timepoint (Figure 5C).

When DSBs persist past MI, the Rad53-dependent DNA damage checkpoint is triggered, causing cells to arrest/delay prior to MII (CARTAGENA-LIROLA *et al*. 2008). The sporulation defect of the +Rap *pif1-md rad54Δ TID1-AA* diploid could therefore be explained if DSBs present at 9 hours triggered this checkpoint after addition of ED promoted passage through MI. Meiotic progression was monitored in the +Rap wild-type and *pif1-md rad54Δ TID1-AA* diploids after addition of ED. The frequency of MI cells peaked at 11 hours in the wild-type strain, while MI cells slowly accumulated and had not peaked after 15 hours in the mutant (Figure 5D). The presence of DSBs results in broken chromosomes so that cells that escape the DNA damage checkpoint and proceed through MII produce fragmented nuclei that can be detected by the presence of >4 DAPI bodies (WENG *et al*. 2024; GAGLIONE *et al*. 2025). While 44% of the wild-type cells progressed through MII, only ∼1% contained fragmented nuclei (Figure 5E and 5F). In contrast, of there was a 3.8 fold increase in the number *pif1-md rad54Δ TID1-AA* post MI cells that contained fragmented nuclei (Figure 5E and 5F). These results further support the hypothesis that *PIF1* functions independently of *RAD54* and *TID1* and is essential for Phase 2 DSB repair when the two translocases are absent.

## DISCUSSION

### A novel assay for studying Phase 2 residual DSB repair during prophase I in yeast

Although there is evidence of a late prophase I transition from interhomolog recombination to intersister recombination in yeast, nematodes and mammals, little is known about the genes and mechanisms that perform this repair (COLAIACOVO *et al*. 2003; HAYASHI *et al*. 2007; ARGUNHAN *et al*. 2017; PRUGAR *et al*. 2017; ENGUITA-MARRUEDO *et al*. 2019; TORAASON *et al*. 2021). This deficit is due in part because of the difficulty in studying Phase 2 recombination due to the low number of DSBs present at pachynema and because new DSBs continue to be made (XU *et al*. 1995; ALLERS AND LICHTEN 2001; THACKER *et al*. 2014; SUBRAMANIAN *et al*. 2016). A new method for Phase 2 recombination was therefore developed that combined reversibly arresting cells in pachynema with preventing new DSB formation by nuclear depletion of Rec104-FRB. This method confirmed that DSB formation and repair both occur continuously during pachynema. When new DSBs were allowed to be made between 7 and 9 hours after transfer to Spo medium, DSBs were still indirectly detected by Hop1 phosphorylation at 9 hours. In contrast, no DSBs were observed 2 hours after Rec104-FRB depletion at 7 hours due to the repair of the “old” breaks. This method was then used to examine Phase 2 recombination in the absence of the candidate genes, *RAD54*, *TID1* and *PIF1*.

### *TID1* can partially substitute for *RAD54* in Phase 2 meiotic DSB repair

The assumption has been that in yeast the transition from Phase 1 to Phase 2 recombination involves changing from Tid1/Dmc1-mediated interhomolog recombination to Rad54/Rad51-mediated intersister recombination (ARBEL *et al*. 1999; NIMONKAR *et al*. 2012; PRUGAR *et al*. 2017). However, whether Rad54 only functions with Rad51 during Phase 2 recombination is unclear. If this simple case were true, then the spore viability of a *rad51-II3A* mutant should be the same as a *rad54Δ*, but instead *rad54Δ* spore viability is considerably worse (Figure 2E)(SHINOHARA *et al*. 1997; CLOUD *et al*. 2012). The more severe *rad54Δ* phenotype could be because Rad54 also can also work with Dmc1 when the Rad51 recombinase is unable to mediate strand invasion, or because Rad54 has other functions besides stimulating D-loop formation such as helping with the homology search (CRICKARD 2021).

While Rad54 may be the preferred translocase for Phase 2 recombination in a wild-type meiosis, robust repair was observed in *rad54Δ* using our method, thus it is not essential. Depletion of Tid1 during pachynema does not increase the appearance of DSBs indicating Tid1 is not normally involved in Phase 2 recombination (SUBRAMANIAN *et al*. 2016). However, when Tid1 was depleted from the *NDT80-IN REC104-AA rad54Δ* diploid, Hop1 phosphorylation increased and spore viability went down compared to *rad54Δ* alone. Therefore, Tid1 can partially compensate for the absence of *RAD54* in residual DSB repair. Although Rad54 and Tid1 bind to different places on the presynaptic filament, there is previous evidence of crosstalk between these two translocases, as a *rad54Δ tid1Δ* diploid exhibits synergistic defects in interhomolog recombination and spore viability (Figure 4C)(SHINOHARA *et al*. 1997; CRICKARD *et al*. 2020). Tid1 and Rad54 can stimulate both Dmc1 and Rad51-mediated D-loop activity *in vitro* (CHI *et al*. 2006; CHI *et al*. 2009; NIMONKAR *et al*. 2012; BUSYGINA *et al*. 2013; SANTA MARIA *et al*. 2013). Whether Tid1 in the *rad54Δ* mutant functions with Rad51 and/or Dmc1 for Phase 2 recombination remains to be determined.

### *PIF1* functions independently of *RAD54* and *TID1* in Phase 2 meiotic DSB repair

Rad51 strand exchange activity is normally inhibited during Phase 1 meiotic recombination by Mek1-dependent mechanisms that prevent interaction with Rad54. However, when this regulation is abolished in a *hedΔR* diploid, the frequency of interhomolog recombination is nearly the same as wild type, with just a two-fold decrease in interhomolog bias (LAO *et al*. 2013; LIU *et al*. 2014; ZIESEL *et al*. 2022). These Rad51-mediated crossovers differ from Dmc1-dependent crossovers in wild-type cells in two ways. First, DSB repair takes longer and requires time provided by the meiotic recombination checkpoint (MRC). Elimination of this checkpoint in *hedΔR* strains lowers spore viability to an average of ∼60% due to an increase in MI non-disjunction (ZIESEL *et al*. 2022). Second, although Pif1 is inhibited by the Mer3-MutLβ complex in wild-type cells and therefore is not involved in Dmc1-mediated recombination, *PIF1* promotes Rad51-mediated interhomolog crossover formation, spore viability and proper MI chromosome segregation during Phase 1 in the *hedΔR* background (VERNEKAR *et al*. 2021; ZIESEL *et al*. 2022).This function requires Pif1 helicase activity and interaction with PCNA. Furthermore, there is a dramatic decrease in spore viability when *PIF1* is depleted from *dmc1Δ hedΔR* diploids where Rad51 is the only recombinase, and intersister DSB repair is delayed when Mek1-as is inactivated in a *dmc1Δ mek1-as pif1-md* strain (Figure 1)(ZIESEL *et al*. 2022). These results led to the idea that Pif1 helps extend invading strands in Rad51-mediated D-loops when Rad51 is constitutively active, similar to its role in break induced replication (WILSON *et al*. 2013; BUZOVETSKY *et al*. 2017; ZIESEL *et al*. 2022).

Constitutive activation of Rad51 during Phase 1 is an artificial situation that does not occur in wild-type meiosis. In contrast, Phase 2 recombination occurs when Mek1 levels have decreased due to synapsis leading to Rad51 activation. We therefore hypothesized that the biologically relevant role for Pif1 is in promoting Rad51-mediated DSB repair during Phase 2. In fact, co-induction of *PIF1* with *NDT80* rescued the modest decrease in spore viability observed in *pif1-md*. This result demonstrated that *PIF1* functions after Phase 1 recombination is complete.

The simple idea that Pif1 extends Rad51-mediated D-loops during Phase 2 recombination was not supported, however, by epistasis analyses showing synergistic spore viability defects when *pif1-md* was combined with either *rad54Δ* or *rad51-II3A*. Further evidence that *PIF1* has a different role in Phase 2 recombination from that observed in the Phase 1 recombination in the *hedΔR NDT80-mid* strain was obtained using the Phase 2 recombination method. In the *pif1-md* mutant alone, no improvement in spore viability was observed when Rec104 was depleted, indicating that a subset of cells had DSBs that were not repairable by using either *RAD54* or *TID1*. In addition, while *PIF1* was not required for the *TID1*-dependent DSB repair that occurs in *rad54Δ,* it became essential in the absence of both *RAD54* and *TID1*.

We conclude that *PIF1* normally has a minor role in Phase 2 recombination but serves as a backup when the translocases are missing. Rad54 has been shown to convert nascent D-loops into mature D-loops by sequentially removing Rad51 monomers from the invading so that complementary base pairing between the invading and donor strands can form heteroduplex DNA (WRIGHT AND HEYER 2014). Tid1 is also able to remove Rad51 from DNA *in vitro*, as well as Dmc1 *in vivo* (CHI *et al*. 2006; HOLZEN *et al*. 2006). One function of Pif1 is the removal of proteins from DNA, such as the Cdc13 telomere binding protein, the Sub1 transcription factor and proteins that block progression of the replication fork (SCHAUER *et al*. 2020; SPARKS *et al*. 2020; CHIB *et al*. 2023). We propose that in the absence of Rad54 and Tid1, Pif1 may move in a 5’-3 direction to remove Rad51 and Dmc1 from the invading strand to allow heteroduplex formation. Alternatively, a small subset of nascent D-loops may normally be created by presynaptic filaments lacking the translocases and extension of the invading strand from this type of D-loop could require *PIF1*.

In summary, this work has established a novel method for studying Phase 2 meiotic recombination in budding yeast that has revealed functional interactions between *RAD54*, *TID1* and *PIF1*. This assay can now be applied to other recombination genes to get a more wholistic understanding of meiotic residual DSB repair.

## DATA AVAILABILITY

Plasmids, strains and custom antibodies are available upon request. The authors affirm that all data necessary for confirming the conclusions of the article are present with the article, Figures and tables. All of the raw data used for the Figures in this paper can be found in Supplementary Data 1. Statistical analyses were performed using GraphPad Prism 9.0 for Mac.

Supplemental material is available at GENETICS online.

## ACKNOWLEDGEMENTS

We are grateful to Bruce Futcher, Michael Lichten, Ed Luk, Aaron Neiman and members of the Hollingsworth lab for helpful discussions. Aaron Neiman provided comments on the manuscript. Thanks to Bruce Futcher and Bob Gaglione for plasmids, Ting-Fan Wang for the Hop1-pT318 antibody, Andreas Hochwagen, Franz Klein and Andrew Ziesel for strains.

## SUPPLEMENTAL FIGURE LEGENDS

**S1 Figure.** Distribution of viable spores in tetrads in the helicase dead *pif1-K264A* mutant and the *pif1-L354P* mutant. (A) The percent of tetrads with the indicated number of viables spores from the dissection data shown in Figure 2A. The WT and *pif1-md* control data were previously published in (ZIESEL *et al*. 2022). The total number of tetrads for each strain are WT (692), *pif1-md* (332), *NDT80-IN* (310), *NDT80-IN pif1-md* (276), *NDT80-IN pif1-md::PIF1-IN* (596). (B) The percent of tetrads with the indicated number of viables spores from the dissection data shown in Figure 2B. The *NDT80-IN* and *NDT80-IN pif1-md* data are the same as in panel A. The total number of tetrads for the remaining strains are *NDT80-IN pif1-md::pif1-m1-3xFLAG-IN* (373), *NDT80-IN pif1-md::pif1-m1-K264A-3xFLAG-IN* (304), *NDT80-IN pif1-md::pif1-m1-L354P-3xFLAG-IN* (473). (C) Structure of *S. cerevisiae* Pif1 (amino acids 287-780) bound to single stranded DNA (LU *et al*. 2018). DNA is indicated in orange, the α-helix containing the Pif1 Family Specific Sequence (PFSS) is pink, the pin-loop structure is green, L354 and K264 are indicated in red and blue, respectively. (D) Alignment of the human (Hs) and yeast (Sc) Pif1 region that contains the PFSS α-helix (indicated by the pink box). Identical amino acids are blue. The cognate L319 and L354 amino acids from human and yeast Pif1, respectively, are highlighted in yellow.

**S2 Figure. Quantification of Hop1-pT318 in various *NDT80-IN REC104-AA* diploids.**

Protein gels from two biological replicates for the strains shown in Figure 5C were probed with Hop1-pT318, Hop1 and Arp7 antibodies. The amount of Hop1-p318 was divided by the amount of total Hop1 and these values were calculated as the percent of the *NDT80-IN REC104-AA* wild-type strain.

